# Glucose and NAADP trigger elementary intracellular β-cell Ca^2+^ signals

**DOI:** 10.1101/2020.11.01.363804

**Authors:** Paula Maria Heister, Trevor Powell, Antony Galione

**Author notes:** **Correspondence**, Correspondence should be addressed to Dr Paula Heister or Professor Antony Galione.

## Abstract

Pancreatic β-cells release insulin upon a rise in blood glucose. The precise mechanisms of stimulus-secretion coupling, and its failure in Diabetes Mellitus Type 2, remain to be elucidated. The consensus model, as well as a class of currently prescribed anti-diabetic drugs, are based around the observation that glucose-evoked ATP production in β-cells leads to closure of cell membrane ATP-gated potassium (K_ATP_) channels, plasma membrane depolarisation, Ca^2+^ influx, and finally the exocytosis of insulin granules (Ashcroft et al., 1984; Cook and Hales, 1984). However, it has been demonstrated by the inactivation of this pathway using genetic and pharmacological means that closure of the K_ATP_ channel alone may not be sufficient to explain all β-cell responses to glucose elevation (Henquin, 1998; Seghers et al., 2000). Here we show using total internal reflection fluorescence (TIRF) microscopy (Axelrod, 1981) that glucose as well as the Ca^2+^ mobilising messenger nicotinic acid adenine dinucleotide phosphate (NAADP), known to operate in β-cells (Johnson and Misler, 2002; Masgrau et al., 2003), lead to highly localised elementary intracellular Ca^2+^ signals. These were found to be obscured by measurements of global Ca^2+^ signals and the action of powerful SERCA-based sequestration mechanisms at the endoplasmic reticulum (ER). This is the first demonstration of elemental Ca^2+^ signals in response to NAADP, although they have been suspected (Davis et al., 2020). Optical quantal analysis of these events reveals a unitary event amplitude equivalent to that of known elementary Ca^2+^ signalling events, inositol trisphosphate (IP_3_) receptor mediated blips (Parker et al., 1996; Parker and Ivorra, 1990), and ryanodine receptor mediated sparks (Cheng et al., 1993). We propose that a mechanism based on these highly localised intracellular Ca^2+^ signalling events mediated by NAADP may initially operate in β-cells when they respond to elevations in blood glucose.

---

The idea that stimulus secretion coupling involves mechanisms in addition to the K_ATP_ channel-mediated pathway is not new. Possible mechanisms include (1) an amplification of the K_ATP_ channel dependent pathway which remains functionally silent until the latter has depolarised the membrane (Henquin, 2000), (2) an additional triggering pathway that converges with the K_ATP_ channel mediated pathway on membrane depolarisation, and (3) an independently functional pathway that may lead to insulin release in the absence of K_ATP_ channel involvement (Islam, 2010). One potential trigger of stimulus-secretion coupling, which may be synergistic with, or independent of, K_ATP_ channel closure, is the local release of Ca^2+^ from intracellular stores (Arredouani et al., 2015). Indeed, there has been some debate about the relative importance of extracellular Ca^2+^ influx versus release from intracellular stores during biphasic insulin secretion, with some proposing that for the first phase of insulin release, influx is dispensable (Wollheim et al., 1978). In terms of subcellular localized Ca^2+^ signals, it was shown in the late 1970s using pyroantimonate precipitation that incubation of β-cells with high glucose led to a Ca^2+^ increase immediately beneath the cell membrane (Wollheim and Sharp, 1981). The possible role of small lysosomal Ca^2+^ stores in stimulus-secretion coupling (Arredouani et al., 2015) warranted the investigation of Ca^2+^ signalling in β-cells at high spatial and temporal resolution. Thus the present study sought to characterise sub-membrane Ca^2+^ transients observed in β-cells loaded with fluo-4 in response to glucose and the membrane permeable form of the lysosomal Ca^2+^ mobilizing messenger, NAADP (NAADP-AM) (Parkesh et al., 2008), using total internal reflection fluorescence (TIRF) microscopy. Furthermore, we hypothesized that dissection of these Ca^2+^ transients under experimental conditions where globalized Ca^2+^ signals or ER-based Ca^2+^ sequestration (Worley et al., 1994) were avoided, might reveal their substructure, akin to the sparks and puffs observed for IP_3_ and ryanodine receptors, respectively (Cheng et al., 1993; Parker et al., 1996; Parker and Ivorra, 1990).

A feature of Ca^2+^ signalling dynamics in pancreatic β-cells is their heterogeneity upon stimulation with the recent proposal that some cells in the islet act as Ca^2+^ signalling hubs or pacemakers (Rutter et al., 2017), whilst others follow through gap-junctional or paracrine signalling mechanisms. However, heterogeneity is also seen in isolated cells. Indeed, Ca^2+^ responses to glucose in β-cells have been described as having a typical triphasic shape often seen in parallel with, and assumed to be the result of, simultaneously measured changes in membrane potential, which have a similar pattern and are in synchrony with the cytosolic Ca^2+^ increase (Worley et al., 1994): Phase 0 consists of a ‘dip’, or initial decrease, in cytosolic free-Ca^2+^ resulting from increased sarco-/endoplasmic reticulum Ca^2+^ ATPase (SERCA) activity transporting Ca^2+^ into the endoplasmic reticulum (ER) in response to the rising ATP concentration following glucose metabolism. Phase 1 constitutes a transient rise in Ca^2+^ associated with L-type Ca^2+^ channel activation and Ca^2+^-induced Ca^2+^ release (CICR) from intracellular stores. Phase 2 are Ca^2+^ oscillations superimposed on a steadily elevated plateau thought to be the result of Ca^2+^ influx through L-type channels. While this standardised model is useful; it has been demonstrated that β-cells display more complex responses to glucose and other stimuli such as GLP1, insulin, and nutrients including amino acids (Gustavsson et al., 2006; Herchuelz et al., 1991). A classification of primary human β-cell autocrine Ca^2+^ responses to insulin shows that β-cells display a variety of equally common but different Ca^2+^ signals and no clear ‘standard’ response (Johnson and Misler, 2002).

We report here sub-membrane Ca^2+^ transients evoked by 16.5 mM glucose in mouse primary pancreatic β-cells, measured using evanescent-wave TIRF microscopy. Recordings were made from 1,017 cells. Fig. 1 shows the classification of typical responses of 239 cells from male WT mice of one genetic background, to avoid additional confounding factors. There was a clear heterogeneity of responses. While over 96% of cells exhibited a clear elevation of sub-membrane Ca^2+^ (calculated as ΔF/F_0_, where ΔF is the change in fluorescence intensity from prestimulation F_0_) ~60% resembled the standardised triphasic profile, with the remaining responding cells described by 3 further classifications (cf. Fig. 1). These data are consistent with the notion that there are β-cells with different patterns of expression of Ca^2+^ channels, which may serve multiple functions within the islet (Rutter et al., 2017). Treatment of mouse primary pancreatic β-cells with extracellular NAADP-AM (10 μM) resulted in similar Ca^2+^ responses to those observed with glucose described above. Ca^2+^ oscillations could be resolved that culminated in a raised plateau of elevated [Ca^2+^]_I_ (Fig. S1).

**Figure 1.**
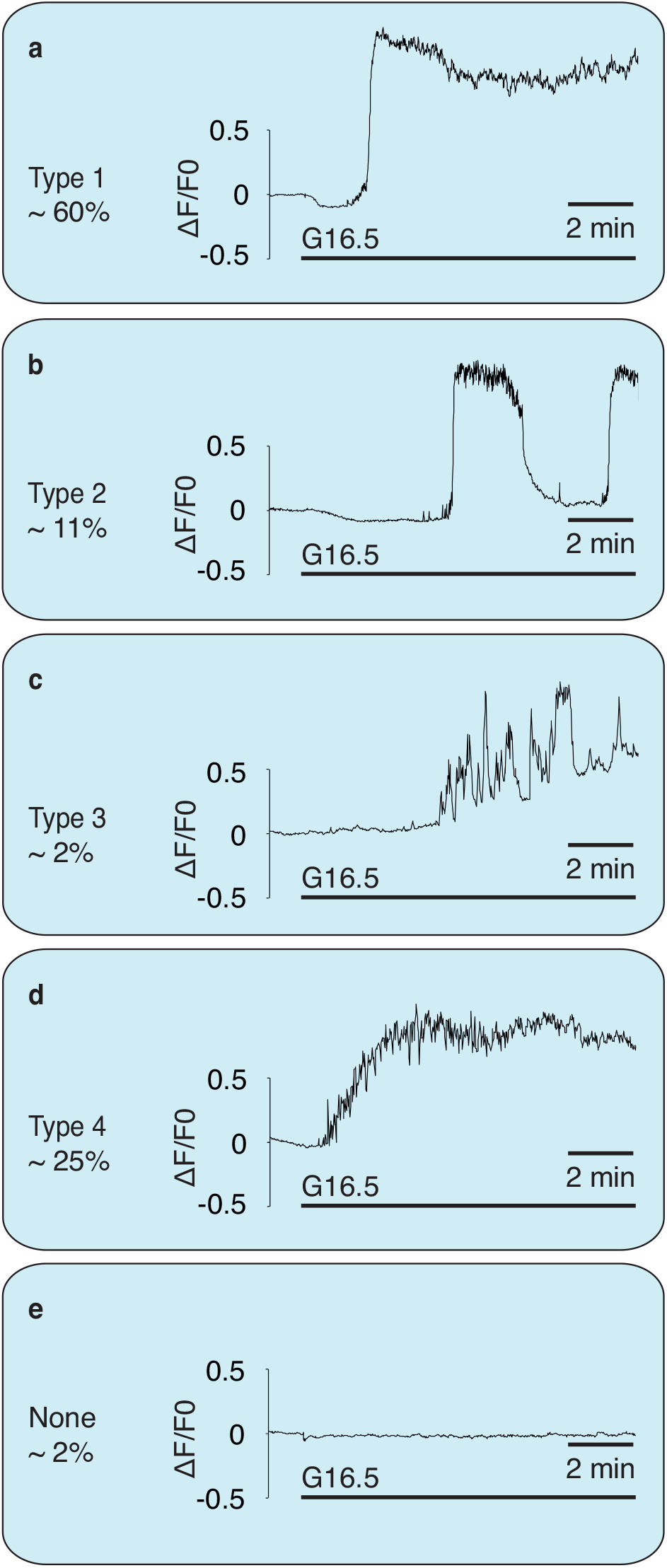
Classification of β-cell sub-membrane Ca^2+^ responses to elevation in glucose as recorded with TIRF. Responses to 16.5 mM glucose at 37° C in mouse primary pancreatic β-cells. n = 239. Responses from 1,017 cells were analysed overall. Percentages are of 239 control cells of male WT mice of one genetic background to exclude potential genetic- or sex-differences in response type distribution. Approximately 96 % showed a prominent global Ca^2+^ response. In 116 cells, 16.5 mM glucose evoked a mean peak height of 2.4 ± 0.01 ΔF/F_0_. (**a**) The most common response (type 1, ~ 60 % of responses) resembles the standardised triphasic response. (**b**) The slow Ca^2+^ oscillations classified as type 2 often occurred at periods of ~ 5 min. **(c)** Response type 3 consists of numbers of large transients superimposed on a slow increase in Ca^2+^. These responses were rare (~ 2 %). **(d)** Type 4 responses (~ 25 %) resemble the triphasic response, but with a less steep Ca^2+^ rise. **(e)** A very small number of cells did not respond to glucose (~ 2 %).

If Ca^2+^ release from sub-membrane stores triggered the global Ca^2+^ response, an increase in Ca^2+^ could be expected to occur first in the vicinity of the cell membrane, the location of the cell’s secretory vesicles which form part of the acidic organelle continuum in the β-cell (Orci et al., 1986), before recruiting a global Ca^2+^ response (Arredouani et al., 2010). Lysotracker was utilised to confirm the presence of acidic stores within the TIRF plane (see Fig. S2). In parallel recordings using TIRF and standard epifluorescence (to monitor global Ca^2+^), however, the two transients were largely superimposable (see supplementary data, Fig. S3). Thus if there were sub-membrane Ca^2+^ release events preceding the global Ca^2+^ response, they were either very rapid and too small to be detected by the current protocol, masked by concomitant larger L-type Ca^2+^ channel-mediated influx, or obscured by increased SERCA pump activity following enhanced glucose metabolism to ATP.

In order to minimise these possible confounding factors, cells were preincubated in recording medium containing 1.7 mM Ca^2+^, 3 mM glucose, and 1 μM of the irreversible SERCA inhibitor, thapsigargin. Immediately before recording, the extracellular fluid was exchanged for Ca^2+^-free medium, containing thapsigargin and either 100 μM or 5mM EGTA. The removal of Ca^2+^ immediately before recording prevented non-ER stores, which may rely on Ca^2+^ influx and ER Ca^2+^ transfer for filling (Morgan et al., 2013; Xu and Ren, 2015), from run-down during the incubation period. Cells were then challenged with either 100 nM NAADP-AM or 16.5 mM extracellular glucose. Over the ~10 min period following either challenge, cells showed clear very brief sub-membrane Ca^2+^ transients with maximum amplitudes >2ΔF/F_0_. These show comparable kinetics to those of Ca^2+^ puffs evoked by IP_3_ (Smith and Parker, 2009). The movie (supplementary information – movie 1) illustrates recordings of these events and examples are shown in Fig. 2a – i. Their localised nature is illustrated in the maximum ΔF/F_0_ trace (Fig. 2m) and the 3D fluorescence intensity plots shown in Fig. 2 j – l. Similar localized Ca^2+^ signals were observed in extracellular solutions containing 100 μM and 5 mM EGTA (cf Fig. S4), strongly suggesting that Ca^2+^ release from intracellular stores but not Ca^2+^ influx are involved in the generation of these elementary Ca^2+^ events. These local transients were also recorded at a higher acquisition rate (46 Hz) (Fig. S5), giving additional resolution of the events captured at our standard experimental recording rates of 3.3 Hz.

**Figure 2.**
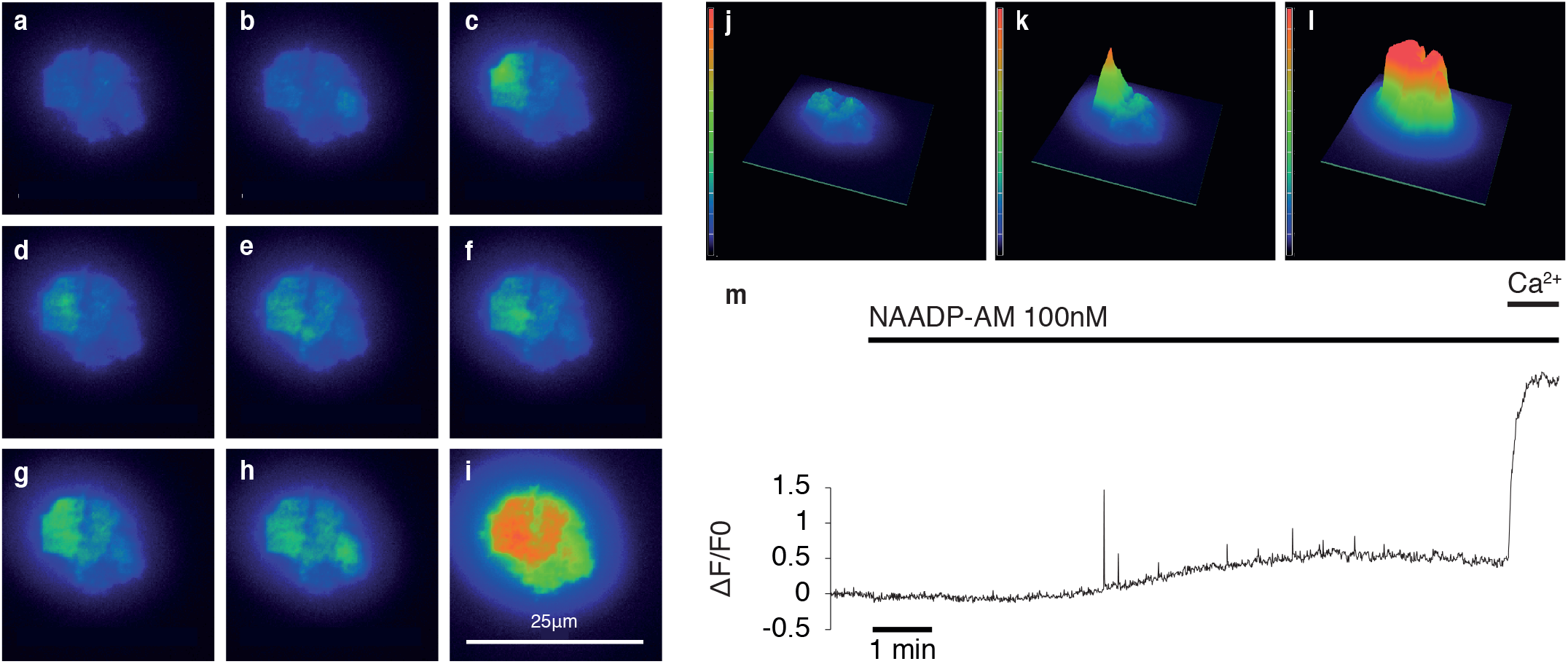
Localised and global Ca^2+^ responses. Responses of a β-cell cluster (3 cells) after NAADP-AM addition as observed using TIRF microscopy in the presence of 5 mM EGTA. **(a)** Baseline fluorescence. **(b – h)** Localized Ca^2+^ release events (for illustrative purposes, larger events were selected). **(i)** Global Ca^2+^ influx. **(j-l)** Intensity profiles of a β-cell cluster, **(j)** at baseline, **(k)** in the presence of an elementary event after addition of NAADP-AM, **(l)** during global Ca^2+^ influx. Images are pseudo-coloured with warmer colours representing higher levels of fluorescence. **(m)** Representative trace of a β-cell showing spiking Ca^2+^ events triggered by NAADP-AM. Maximum intensity change after normalisation to baseline is plotted against time. Extracellular Ca^2+^ was re-admitted at the end of the experiment; leading to a global Ca^2+^ response.

To resolve these individual events in more detail, a “spark” detection algorithm was used (see Methods) which also removed the underlying ‘ramp’ in cell Ca^2+^ that can be seen in Fig. 2m. Clearly, there is an increasing frequency of events following application of 100 nM NAADP-AM or 16.5 mM glucose to a maximum; followed by a gradual decline (Fig. 3 a, b). This is in accordance with the self-limiting nature of these types of signals when stores cannot be replenished during sustained stimulation (Cheng and Lederer, 2008) or desensitization to signals such as NAADP (Cancela et al., 1999; Johnson and Misler, 2002). To quantify the changes in frequency, the mean percentage of frames showing at least one spike (defined as an event 2 standard deviations above mean baseline ΔF/F_0_) was determined for the baseline (last 80-100 frames before stimulus) and after the addition of raised glucose or NAADP-AM. As illustrated in Fig. 3e and f, both 100 nM NAADP-AM and 16.5 mM glucose evoked an approximately ten-fold increase in spike frequency.

**Figure 3.**
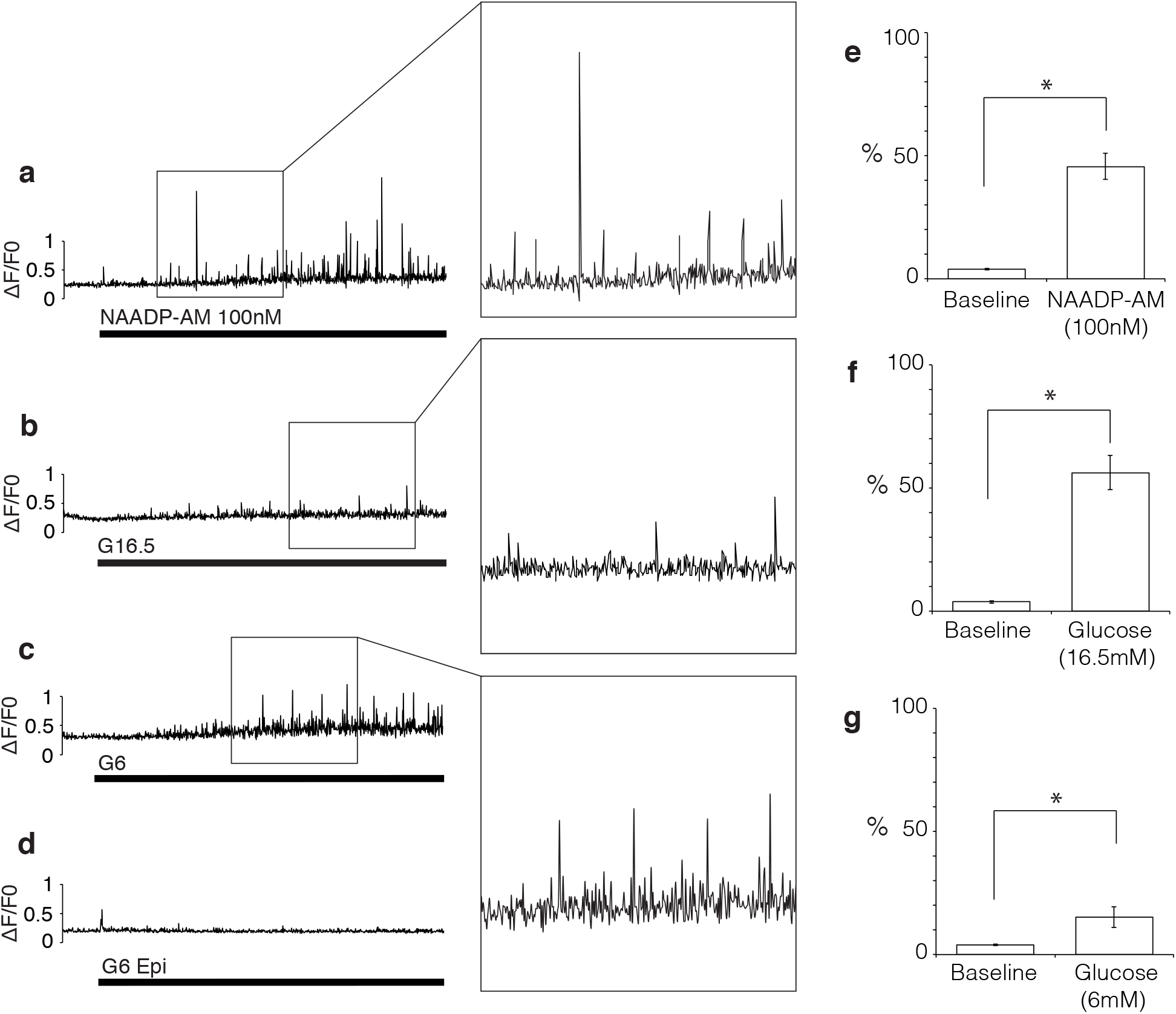
Quantification of calcium release events in response to glucose and NAADP-AM in low EGTA (100μM). **(a – c)** Representative TIRF traces of β-cells stimulated with: **(a)** 100 nM NAADP-AM, **(b)** 16.5mM glucose, **(c)** 6 mM glucose. **(d)** Epifluorescence recording (all other parameters equal) using 6 mM glucose. Maximum intensity change of subsequent frames after normalising to baseline plotted against time. Insets show magnified events for each of the three experimental stimuli, chosen to illustrate the variation in event size. **(e – g)** Percentage of frames showing elementary events (defined as events of an amplitude more than 2 standard deviations above baseline mean) before (baseline) and after stimulus addition. **(e)** NAADP-AM; n=20 (8 experiments, 4 Animals), t(19)=7.90, p < 0.01. **f** 16.5 mM glucose; n = 14 (6 experiments, 4 Animals), t(13)=7.44, p < 0.01. **g** 6 mM glucose; n = 18 (6 experiments, 6 Animals), t(17)=2.61, p < 0.01. * denotes significance; paired samples, one-tailed t-tests.

Having unmasked discrete Ca^2+^ signals in response to a high concentration of glucose (16.5 mM), this raises the question of whether similar responses could be evoked by a smaller increase in extracellular glucose. We examined whether a doubling of glucose from 3 mM to 6 mM (a stimulatory concentration within the physiological range) without thapsigargin pre-incubation, would still allow us to detect visible Ca^2+^ events with TIRF. We reasoned that at lower concentrations of glucose, less ATP would be produced resulting in lower SERCA activity, which would otherwise obscure detection of local Ca^2+^ events. As shown in Fig. 3c, the effects of stimulation with low glucose in the absence of thapsigargin, resemble those of stimulation with high glucose in the presence of thapsigargin. The percentage of frames containing spikes is significantly higher after raising extracellular glucose to 6 mM (15%) than at baseline (4%; Fig. 3g). Using epifluorescence microscopy, no local Ca^2+^ release events were observed (Fig. 3d).

To analyse the nature of the elementary Ca^2+^ events, optical quantal analysis was carried out on images like those shown in Fig. 4 d-e. Plotting the frame-maxima across all cells stimulated with NAADP-AM results in the frequency distribution depicted in (Fig. 4f). Modes are visible in the ‘tail’ of this decaying function which occur at a period of around 0.05 ΔF/F_0_ (see figure inset). This order of magnitude is comparable to optically-assessed Ca^2+^ blips through IP_3_ receptors (0.1 ΔF/F_0_; (Smith and Parker, 2009) and the smallest imaged IP_3_-evoked Ca^2+^ signals in pancreatic acinar cells (<0.1ΔF/F_0_; (Fogarty et al., 2000). To demonstrate the quantal nature of the Ca^2+^ responses at the level of the individual cell, responses from a representative cell cluster are shown in Fig. 4g. Specific regional areas of interest (ROIs) within the cell were chosen by hand on the basis of their containing at least one Ca^2+^ signalling event. These ROIs were then analysed for their mean amplitudes after subtraction of baseline fluorescence. Events were included if they lay above baseline mean + 2 SDs) of that particular cell. The frequency distribution displays modal behaviour as illustrated by the poly-Gaussian function fitted to it, and the putative unitary event amplitude for this cell cluster is 0.03 ΔF/F_0_.

**Figure 4.**
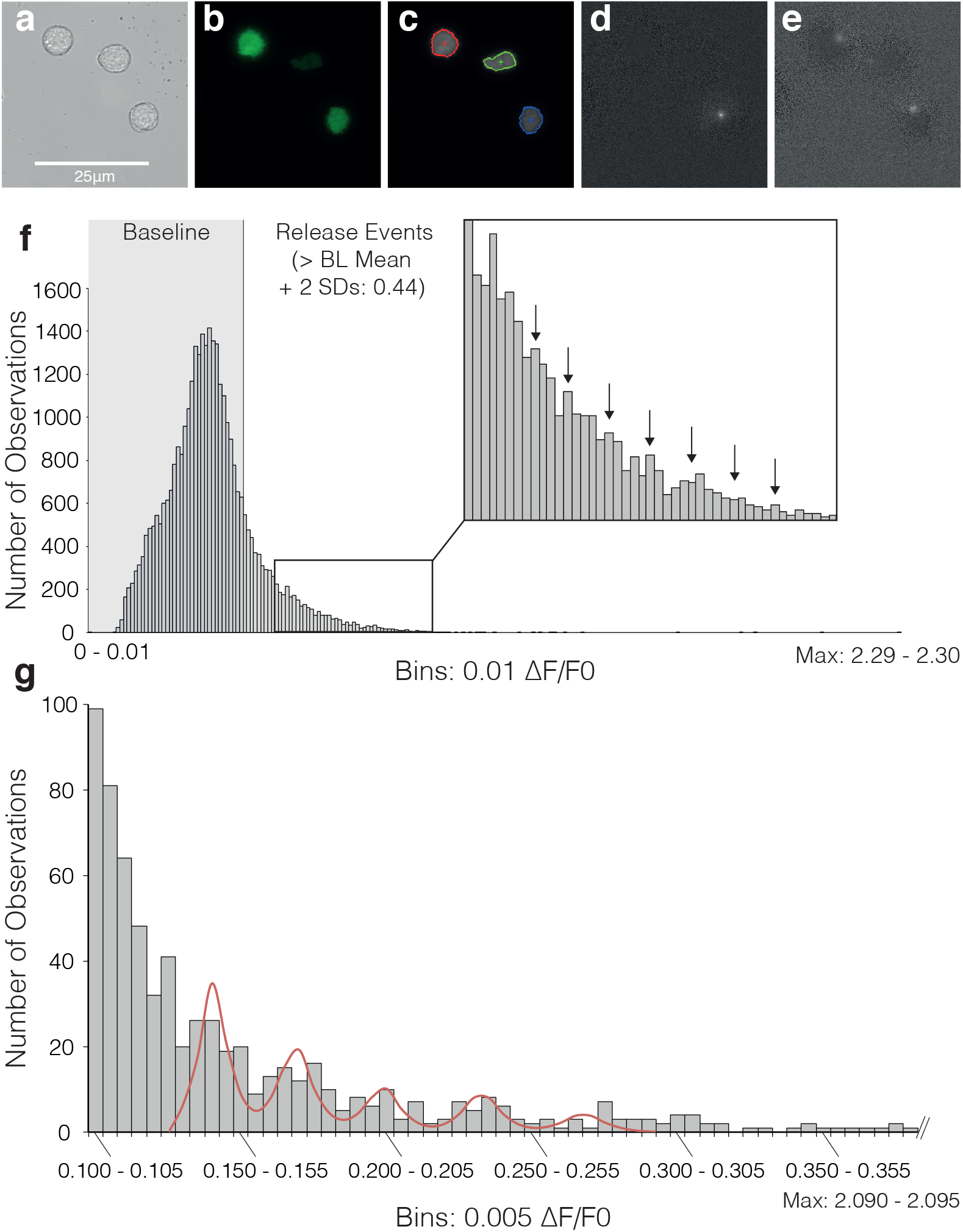
Quantal analysis of elementary events. **(a – e)** Illustration of spark analysis, including examples of elementary release events in response to 16.5 mM glucose detected by normalisation, frame by frame and background subtraction in 3 individual β-cells: **(a)** Original brightfield, **(b)** TIRF, **(c)** Boundaries used in detection analysis, **(d – e)** Individual events isolated by analysis. Note that images of glucose-induced events were chosen for illustration of the analysis mechanism; quantal analysis was carried out exclusively on NAADP-AM stimulated cells. **(f)** Frequency histogram of maximum intensity change > 2 SDs above baseline mean after NAADP-AM Stimulation (n = 20). **(g)** Frequency histogram of mean event amplitudes across areas of interest (n = 51) within a single cell cluster stimulated with NAADP-AM in the presence of 5 mM EGTA. The five peaks of the putative modal distribution in the data were selected as means for individual Gaussian distributions; and 3 to 4 bins to their left and right used to calculate the respective distribution’s standard deviation. The resultant means and standard deviations were used to fit a quintuple Poly-Gaussian function (Sum of five Gaussian functions with Means, SDs: 0.143, 0.007; 0.171, 0.008; 0.201, 0.0089; 0.235, 0.0092; 0.270, 0.0103, respectively) to the data. Histogram was cropped for better resolution-actual maximum intensity bin is 2.09 – 2.095.

The above experiments demonstrate that β-cells show great variability in their Ca^2+^ responses and a high level of spontaneous activity; supporting the notion of a β-cell’s ‘Ca^2+^ fingerprint’ (Prentki et al., 1988). As β-cells are frequently examined in artificially amplified resting and active states during experiments (Islam, 2010, 2020), we have shown here that stimulation with the high glucose concentrations often employed, leading to high SERCA activity (Roe et al., 1994), obscures more subtle Ca^2+^ changes taking place within the cell. It is probable that multiple stores and channels play a part in this process (Galione, 2019; Islam, 2020). The fact that the events occur after pre-incubation with thapsigargin, ruling out the ER as a source, are most striking when triggered with NAADP-AM which targets acidic stores, and are localised just beneath the membrane (the primary location of insulin granules, a subset of the acidic organelles in the β-cell (Orci, 1985)), is a strong suggestion that the source of these events are acidic Ca^2+^ storage organelles. With the discovery of two pore channels (TPCs) as the putative target for NAADP (Calcraft et al., 2009; Ruas et al., 2015), and the present discovery of elementary Ca^2+^ release events in response to NAADP; it is likely that NAADP-evoked Ca^2+^ signals from acidic stores are built from elementary events via activation of TPCs. It was suggested in 1996 that NAADP-mediated Ca^2+^ release was quantal on the basis of its graded release from sea urchin homogenate (Genazzani et al., 1996) and in intact sea urchin eggs localized Ca^2+^ responses ascribed to the osmotic lysis of acidic stores by GPN (Churchill et al., 2002); 25 years on, we may have evidence in a mammalian cell type that a similar principle operates. Thus with the existing sets of elementary Ca^2+^ signals: IP_3_, the IP_3_R, and blips; cADPR, the RyR, and quarks, respectively, elementary Ca^2+^ release events appear to be a governing principle of intracellular Ca^2+^ signalling, regardless of channels involved or Ca^2+^ storage organelle.

In addition to demonstrating elementary Ca^2+^ signals in response to NAADP for the first time, the present study also suggests a potential role for these events in stimulus-secretion coupling in β-cells. NAADP has been shown to elicit Ca^2+^ signals and insulin release in mouse pancreatic β-cells (Arredouani et al., 2015; Johnson and Misler, 2002; Masgrau et al., 2003; Park et al., 2013). Whilst there is argument over the identity of NAADP’s target in the beta cell (Arredouani et al., 2015; Cane et al., 2016; Guse and Diercks, 2018), mutations in the two pore channel gene (TPCN2), the potential principal target for NAADP, have been implied in the inheritance of diabetes type 2 in humans (Fan et al., 2016). Localized Ca^2+^ signals from acidic stores have been proposed to cause depolarisation by activating calcium-dependent cation channels in the plasma membrane as we have previously observed (Arredouani et al., 2015; Foster et al., 2018), such as TRPM4 (Arredouani et al., 2015) and TRPM5 (Brixel et al., 2010; Colsoul et al., 2010; Enklaar et al., 2010). NAADP applied through a patch-pipette evoked small oscillatory cation currents which were preceded by small Ca^2+^ transients (Arredouani et al., 2015) which were abolished in cells from *Tpcn2^-/-^* mice. Importantly, elevating glucose concentrations evoked similar cation currents, which along with those evoked by NAADP were inhibited by the NAADP antagonist, Ned-19 (Arredouani et al., 2015). We have now imaged these localized Ca^2+^ events with our TIRF methodologies in the current study. However, local Ca^2+^ release from acidic stores is also known to trigger CICR in many cells, likely involving membrane contact sites with the ER (Patel and Brailoiu, 2012). In yet other cell types, NAADP was shown to induce localized Ca^2+^ release from secretory granules to initiate their own exocytosis (Davis et al., 2012). In pancreatic β cells, ER Ca^2+^ leak and subsequent uptake into mitochondria via the mitochondrial Ca^2+^ uniporter (MCU) complex have been proposed to prime ATP synthesis (Klec et al., 2019). Since it has been suggested that TPCs can be blocked by ATP (Cang et al., 2013), there may be a complex interplay between Ca^2+^ release from acidic stores and the dynamics of ATP concentrations at the subcellular level.

Thus we propose that during stimulus-secretion coupling in β-cells K_ATP_ channel closure induced by a rise in local ATP levels increases membrane resistance allowing small cation currents, activated by localized Ca^2+^ signals (a summation of elementary Ca^2+^ events from intracellular stores and CICR) to initiate depolarization of the plasma membrane. This in turn results in the opening of L-type Ca^2+^ channels whose mediation of larger globalized Ca^2+^ signals triggers insulin granule exocytosis. This model differs from, but contains elements of, each of the three models of a K_ATP_ channel independent pathway previously discussed. The present investigation suggests that glucose initially generates localized Ca^2+^ signalling events based on Ca^2+^ release from non-ER, likely acidic, stores prior to the influx of Ca^2+^ through VGCCs. We propose that some of the elementary trigger events involved are likely to be mediated by NAADP as we have previously suggested (Arredouani et al., 2010; Arredouani et al., 2015).

## Methods

### Primary β-Cell Culture

Mouse pancreatic islets (from C57BL/6, 10-14 week-old males) were isolated as described previously (Ravier and Rutter, 2010). Islets were dispersed into single cells or cell clusters and plated onto poly-L-lysine coated glass coverslips (Menzel) and incubated in cell* dishes at 37° C for 4 – 7 hours before adding cell culture medium (RPMI 1640, -glucose +glutamine, Gibco), supplemented with penstrep (10,000 U/ml penicillin / 10,000 μg/ml streptomycin, Gibco) and 10 % fetal bovine serum (FBS, Gibco) containing 10 mM glucose and incubated for a further 15 – 17 hours before first use. Cells were loaded with 500 nM fluo 4-AM (Invitrogen) for 1 h in the dark, before being washed with imaging buffer (NaCl 130 mM, KCl 5.2 mM, MgCl_2_ 1 mM, HEPES 10 mM, CaCl_2_ 1.7 mM; pH: 7.4, 280 – 340 mOsm / kg) containing 3 mM glucose and left in this for 10 min before start of recording. Experiments were conducted exclusively at 37° C, and with a baseline glucose concentration of 3 mM, simulating physiological conditions. Experiments to demonstrate localised elementary events were conducted on both single cells and small clusters; at an estimated ratio of 50 % cells, 50 % clusters. Clusters lend themselves to TIRF experiments, as they provide a large area positioned in the same focal-plane. Dispersed cells are usually at marginally different focal planes due to the inherent curvature of the glass coverslip. In studies using Ca^2+^-free medium, extracellular Ca^2+^ was re-admitted at the end of the experiment to visualise a global response as a verification of the cell’s viability. In experiments to visualise acidic stores, cells were preincubated for 10-30 min with LysoTracker Red (Invitrogen) at 200nM.

### Imaging

Cells were excited with an Argon-Ion laser (Andor DU-897, 40 mW; Melles Griot) at 488 nm, and images were obtained using a Nikon Evanescent Wave Imaging System; an Inverted Total Internal Reflection Microscope (Nikon Eclipse Ti) equipped with 60 x and 100 x CFI Apochromat TIRF Series oil-immersion lenses. These lenses have a numerical aperture of 1.49, which allowed for a maximal incident angle of 76.87° calculated by α ≥ sin^-1^ (n_2_ / n_1_), n_1_ > n_2_, where n_1_ is the refractive index of the cover glass (1.53), and n_2_ the numerical aperture of the lens (1.49). The exact angle for experimentation was determined using a Bertrand lens and the beam adjusted. All parameters were controlled using NIS Elements AR 4.0 (Nikon). Images were acquired at a rate of 3.3 Hz for single channel recordings (i.e. only TIRF or Epifluorescence) and at a pre-programmed rate for dual channel recordings (Frame Rate Epi: 1Hz. Frame Rate TIRF: 3.3Hz). Movie 2 and the corresponding data in Fig. S5 were acquired using NIS Elements RAM capture mode (acquisition rate ~46Hz). A CCD camera (Andor iXon+) was used to capture emitted fluorescence at 515 – 555 nm (Binning: 1×1, Exposure: 300 ms, Multiplier: 89, Readout Speed: 10 Hz, Conversion Gain: 1x, Dimensions: 512 x 512, 160nm / pixel).

### Analysis

Fluorescence change across each whole cell (selected as an area of interest, ROI) was analysed by first normalising each image (time frame) pixel by pixel with respect to the average fluorescence across the last 80-100 frames before cell stimulation with glucose or NAADP-AM (cells stimulated using glucose were not used for quantal analysis, as it is an unspecific stimulus and may trigger events via multiple pathways). To eliminate the moving baseline (ramp), a frame by frame subtraction function was used. Of these normalised, baseline-controlled images, the maximum intensity within each cell was measured. A similar algorithm has been used for 2D images previously (Banyasz et al., 2007), and while this paper was in preparation, an automated system applying a sophisticated version of it was released (Ellefsen et al., 2014). From the measured maximum intensities for each cell over time, frequency histograms were compiled to depict the quantal nature of the events, where events more than 2 standard deviations above the baseline mean were considered genuine. This is a standard estimate of minimum spark amplitude (Fogarty et al., 2000). All analysis was conducted in NIS Elements AR 4.0 (Nikon) and MS Excel 14 (Microsoft). Figures were prepared in Illustrator CS6 (Adobe). Ca^2+^ traces are of fluorescence change relative to baseline mean fluorescence (ΔF/F_0_). Base-line mean fluorescence (F_0_) was calculated from the fluorescence of the last 80-100 frames before stimulation. Traces are representative, as indicated in figure legends. Ca^2+^ fluorescence images are pseudo-coloured so that changes in colour reflect changes in fluorescence. Warmer colours represent higher levels of fluorescence. Statistical analysis was conducted in MS Excel 14 and SPSS 19 (IBM). Student’s t-tests (paired or unpaired, one- or two-tailed, as applicable) were used to determine the statistical significance of observed effects (p < 0.05, p < 0.01, as stated). Charts illustrating statistical differences between groups depict mean ± standard error of the mean (SEM) unless stated otherwise. Videos for supplementary information were prepared from Nikon .nd2 files and rendered using Premiere Pro CS6 (Adobe).

## Acknowledgements

This work was funded by a Medical Research Council Programme Grant and Wellcome Trust Senior Investigatorship to AG. PMH was supported by a Department of Pharmacology, University of Oxford DPhil Studentship. We would like to thank Professor Patrik Rorsman for support with experiments and Dr Anthony Morgan for comments on the manuscript.

## Author Contributions

AG, TP and PMH designed the experiments. PMH conducted the experiments. AG and TP supervised the project. PMH, TP and AG prepared the manuscript. All authors reviewed the results and approved the final version of the manuscript.

---

**Figure S1.**
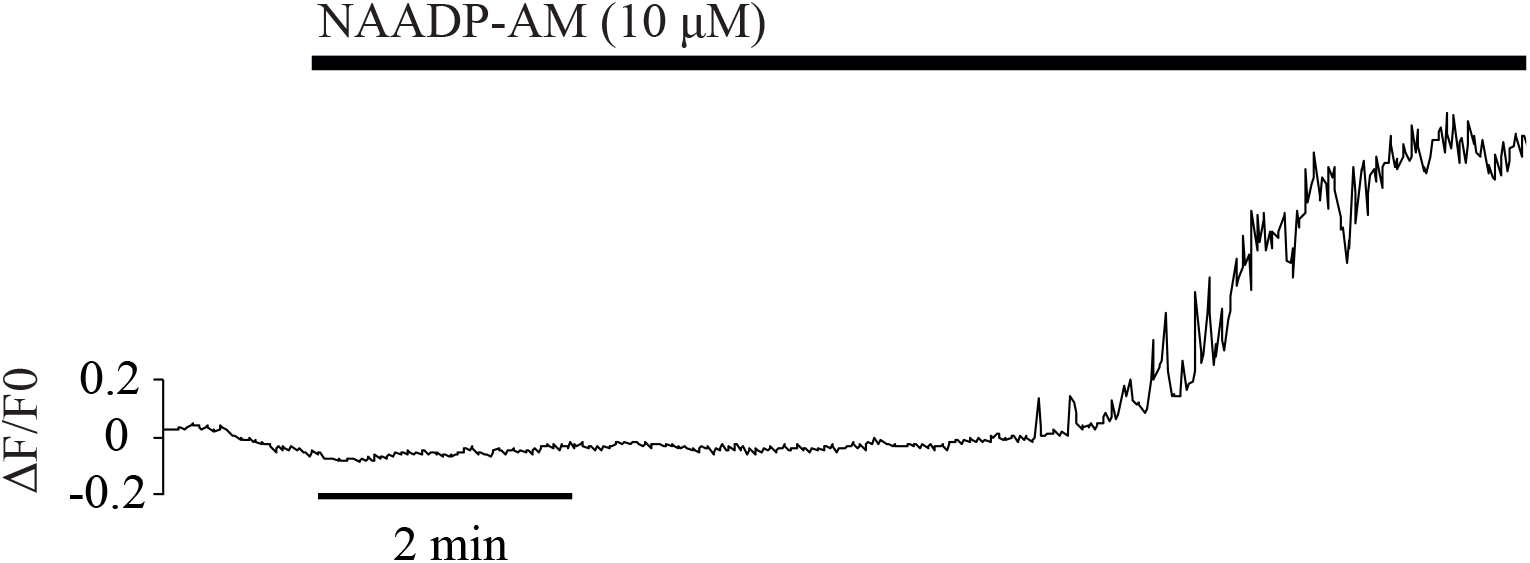
(Extended Data) β-cell sub-membrane Ca^2+^ response to NAADP-AM as recorded with TIRF. Representative trace of a submembrane calcium response to stimulation with 10μM NAADP-AM in the presence of 3mM glucose (used as baseline glucose in all experiments, see methods) imaged using TIRF microscopy.

**Figure S2.**
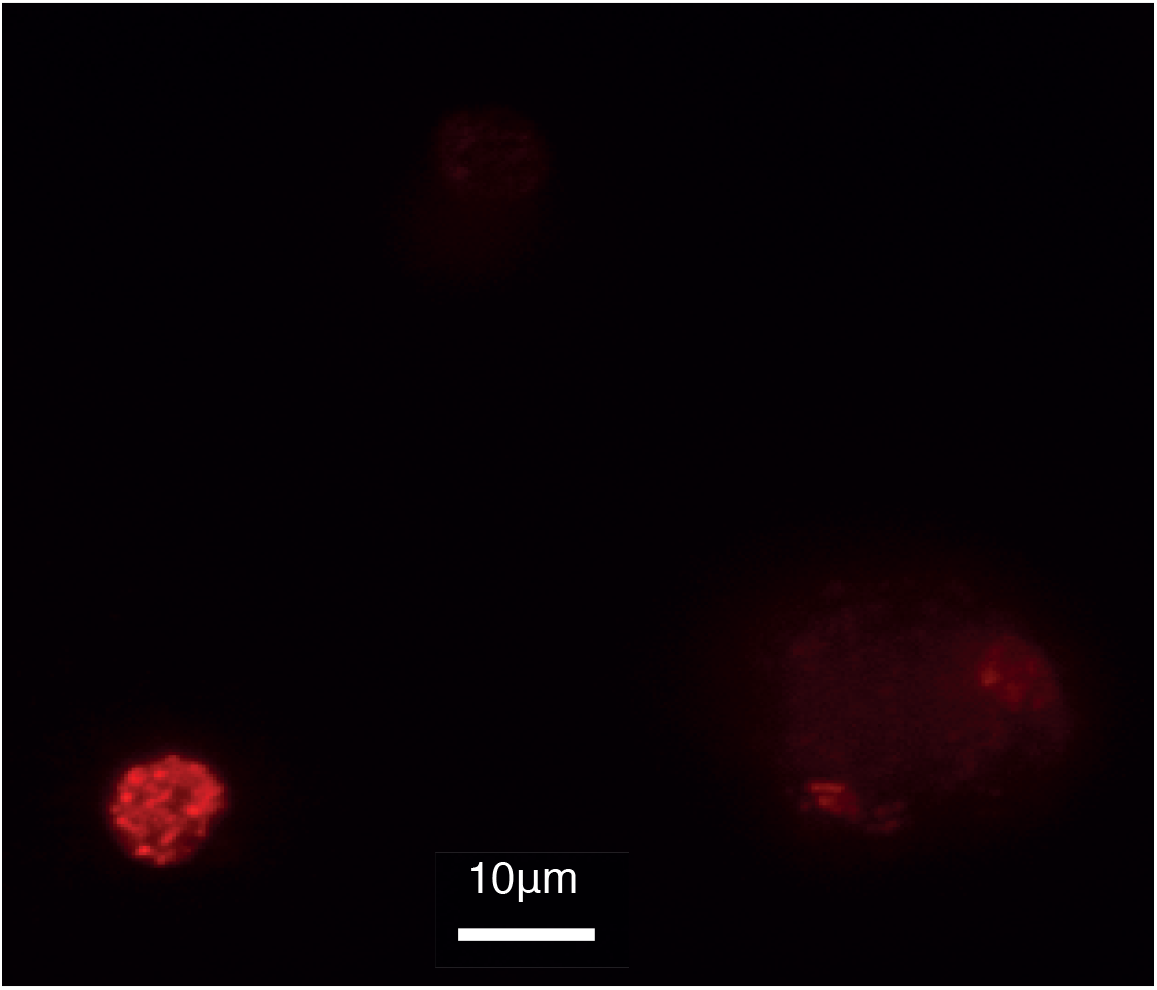
(Extended Data) Visualisation of acidic stores in the TIRF plane. LysoTracker Red labelling of a single β-cell (left) and a β-cells cluster (right) as viewed with TIRF under 60 x magnification.

**Figure S3.**
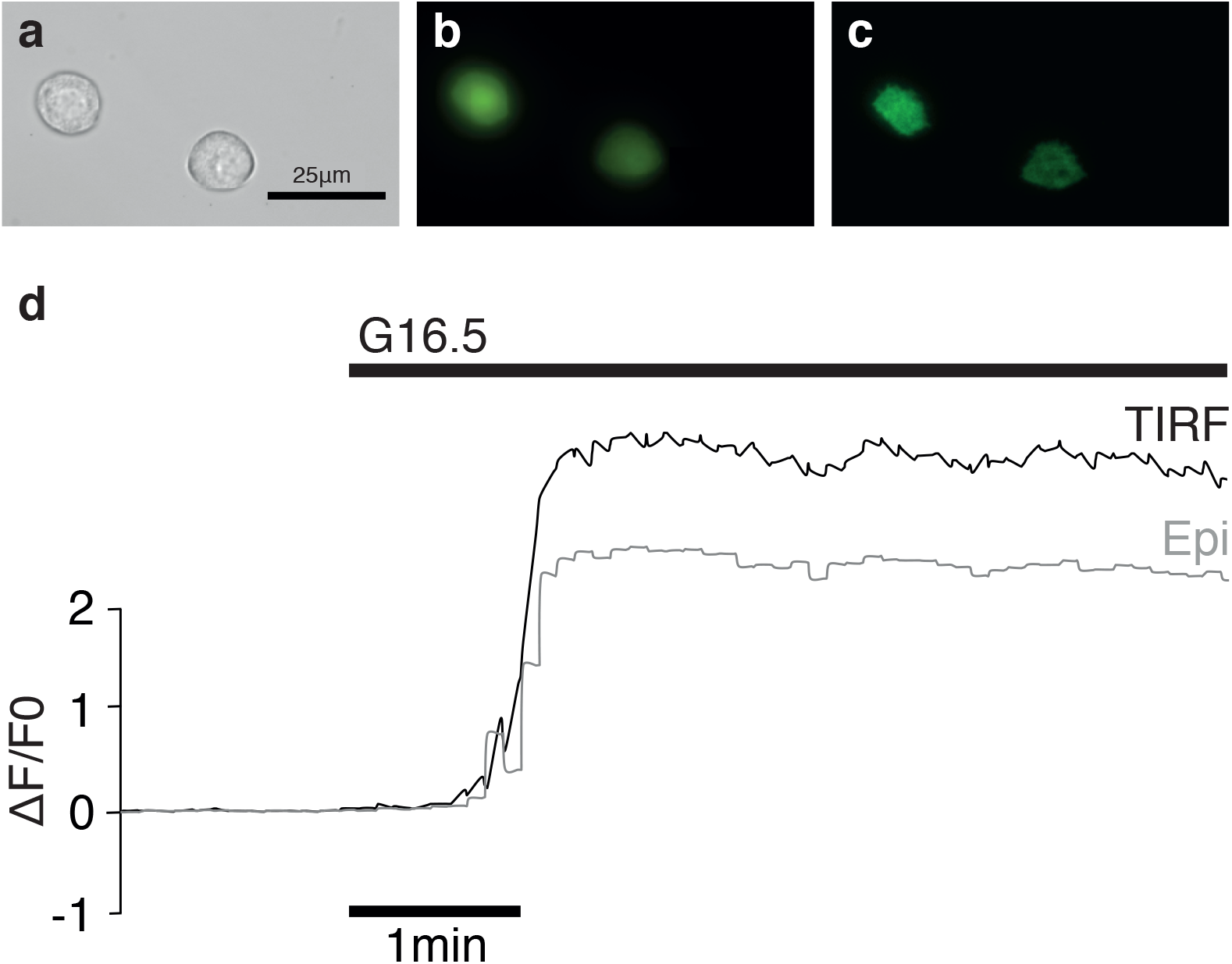
(Extended Data) Parallel imaging of TIRF and epifluorescence in β-cells. Examples of representative **(a)** brightfield, **(b)** epifluorescence, and **(c)** TIRF images of individual β-cells loaded with fluo-4 at 60 x magnification. **d** Representative trace of a parallel recording of whole-cell (epifluorescence; grey trace) and sub-membrane (TIRF; black trace) Ca^2+^ in response to 16.5 mM glucose in a β-cell.

**Figure S4.**
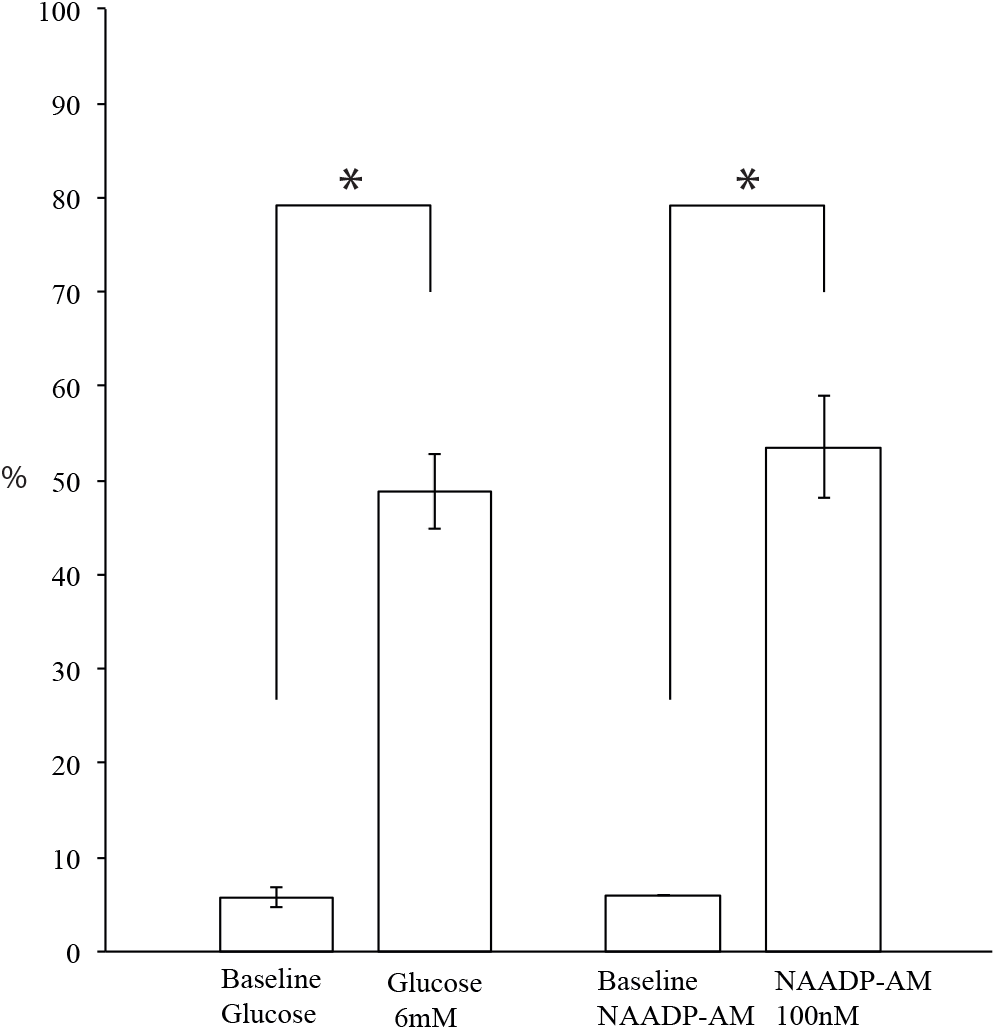
(Extended Data) Quantification of calcium release events in the presence of high EGTA (5mM). Percentage of frames showing elementary events (defined as events of an amplitude more than 2 standard deviations above the baseline mean) before (baseline) and after stimulation with glucose (6mM) or NAADP-AM (100 nM) in the presence of 5mM EGTA. Results are from 4 cells (4 experiments, 2 animals) and 2 cells (2 experiments, 2 animals) respectively. * denotes significance (paired samples, one-tailed Student’s t-test, p<0.01, p<0.05, respectively).

**Figure S5.**
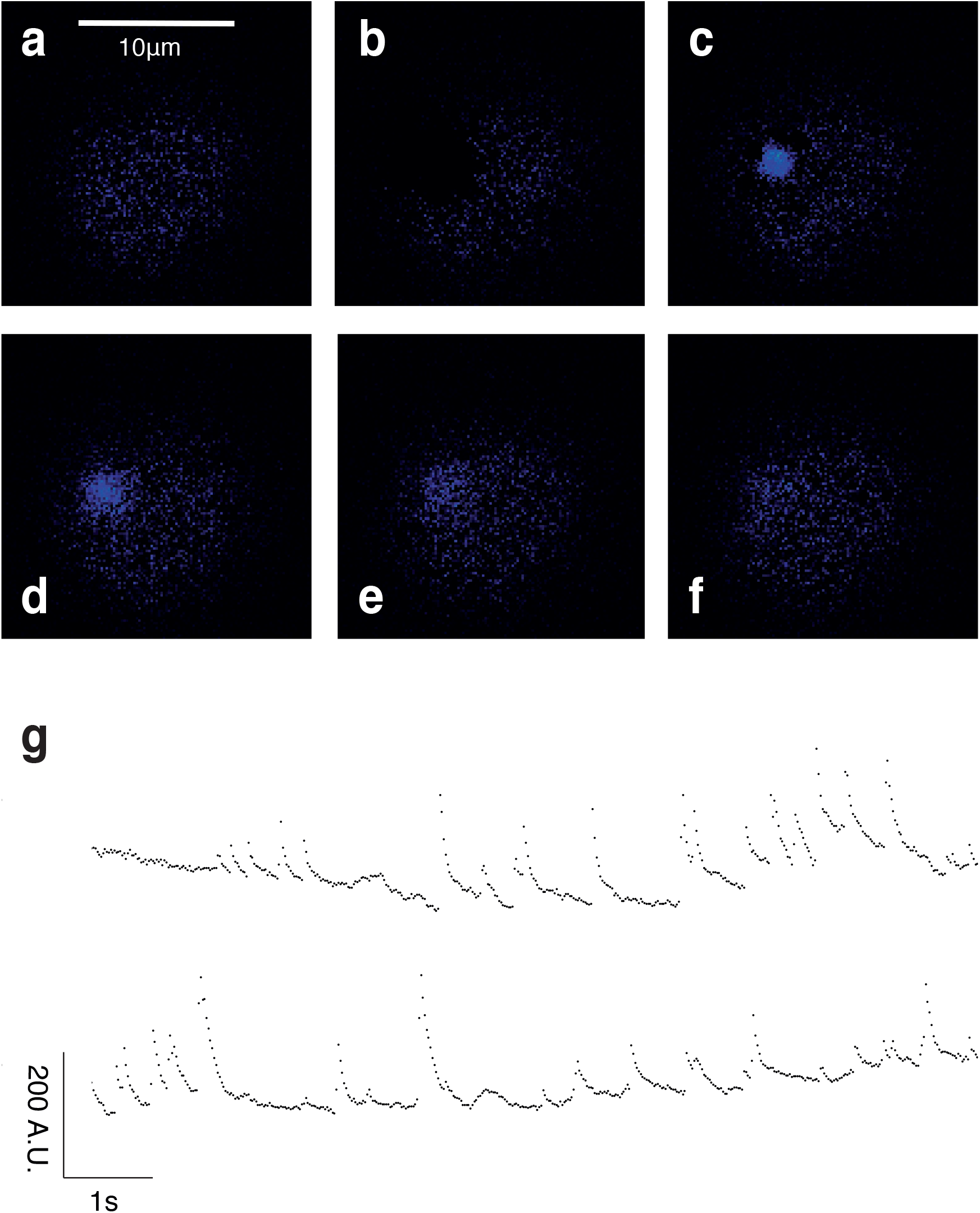
(Extended Data) High speed recordings of calcium events for illustration of diameter and time course. Cells were investigated as above but recorded at ~ 46 frames/second (hardware limit for the setup; proof of method experiment, n = 2, 1 experiment, 1 animal). **a – f** Images of a β-cell over the time course of one individual localised calcium event. **g** fluorescence intensity over time for two individual β-cells (note: traces not normalised, y-axis in arbitrary units, A. U.)

**Movie 1 (Extended Data)** First recording of elementary events in a pancreatic β-cell cluster in response to NAADP-AM (100 nM) visualised using TIRF microscopy at 100 x magnification. At the end of the experiment, extracellular Ca^2+^ is re-admitted, demonstrating a global Ca^2+^ response and cell viability. Recording speed: 3.3Hz, playback speed increased for ease of viewing.

**Movie 2 (Extended Data)** Recording of elementary calcium events in two pancreatic β-cells recorded at RAM capture speed (~ 46Hz) following spark detection (proof of method experiment, n = 2, 1 experiment, 1 animal).

